# Anatomical Identification and Pressure Myography of the Rat Middle Cerebral Artery: A Comprehensive Protocol for Diverse Genetic Models

**DOI:** 10.64898/2026.06.12.718520

**Authors:** Gilbert C. Morgan, Andrew Gregory, Yvan Hanscom-Trofy, Rui Dong, Fan Fan

## Abstract

The middle cerebral artery (MCA) is critical for cerebral blood flow autoregulation and a primary site of cerebrovascular pathology in stroke, Alzheimer’s disease, and vascular dementia. Pressure myography enables precise *ex vivo* quantification of MCA structure and function, but requires accurate anatomical identification and careful vessel handling to ensure reproducibility across diverse rat genetic models. This chapter provides a comprehensive, step-by-step protocol for isolating and cannulating the rat MCA M2 segment for pressure myography. We detail precise anatomical landmarks to ensure consistent vessel selection across strains. The protocol includes optimized solutions, cannulation techniques, and pressure protocols validated across multiple rat models, including transgenic (TgF344-AD), diabetic (T2DN), consomic (SS.5^BN^, FHH.1^BN^), and genome-edited strains. Extensive troubleshooting notes address common technical challenges, including vessel viability assessment, pressure integrity, and strain-specific autoregulatory ranges. This methodology bridges molecular genetic findings with fundamental cerebrovascular physiology, enabling researchers to characterize myogenic reactivity, passive mechanical properties, and structural remodeling in rat models of cerebrovascular disease.

## 1. Introduction

The rat *(Rattus norvegicus)* has historically been one of the premier physiological models for cardiovascular research [1, 2]. Rats share approximately 90% of their DNA with humans, with many large chromosomal regions highly similar, and are well-suited for integrated physiological and pharmacological research due to their manageable size, relatively large blood volume, and stable hemodynamics [3]. Rats develop spontaneous diseases that closely mimic human conditions, most of which involve complex polygenic inheritance. For example, the Spontaneously Hypertensive Rat (SHR) is widely regarded as a “gold standard” model for polygenic essential hypertension [4, 5]. Zucker Rats naturally develop early-onset obesity and insulin resistance, similar to human Type 2 diabetes [6]. The PCK rat carries a spontaneous mutation mimicking human Polycystic Kidney Disease (PKD) [7]. The Fawn-Hooded Hypertensive (FHH) rat spontaneously develops conditions resembling human chronic kidney disease, hypertension, and platelet bleeding disorders [8, 9]. The Dahl Salt-Sensitive (SS) rat is a powerful model that spontaneously mimics human salt-sensitive hypertension, progressive kidney scarring, and the widespread chronic pain and anxiety characteristic of fibromyalgia [10, 11]. Unlike many other models that require obesity (like the Zucker rat), T2DN rats develop spontaneous diabetes and severe insulin resistance without being overweight, which is common in many human Type 2 patients [12–14]. Together, these spontaneous disease models highlight the complexity of polygenic traits, in which multiple genetic variants interact to produce clinical phenotypes, underscoring the need for approaches that can isolate specific genetic contributors.

An important consideration in rat research is the choice between inbred and outbred strains, reflecting a trade-off between genetic homogeneity and heterogeneity [15]. Inbred strains, such as Fischer 344 (F344) or Brown Norway (BN), are produced through at least 20 generations of brother-sister mating, resulting in genetically identical individuals that ensure high experimental reproducibility and a stable background for genetic mapping. In contrast, outbred strains, such as Sprague Dawley (SD) or Wistar rats, maintain a degree of genetic heterozygosity that more closely reflects the diversity of human populations, making them ideal for broader pharmacological screening and general physiological studies. To further dissect polygenic complexity, genome-wide association studies (GWAS) have revealed multiple quantitative trait loci (QTL) associated with complex diseases, and congenic and consomic (chromosome-substitution) rat strains have been used to systematically isolate and dissect these individual polygenic traits [16]. Congenic rats are developed by backcrossing a specific donor genomic segment onto a recipient background, whereas consomic strains involve replacing an entire chromosome. These models enable precise genetic dissection by directly linking physiological phenotypes to specific genomic regions. Notable examples include the FHH.1^BN^ strain [9], used to isolate genetic drivers of hypertension and renal failure, and SS.5^BN^ [10], employed to study the mechanisms of salt-sensitive hypertension. By simplifying complex systemic diseases into single-locus analyses, these strains provide a robust platform for investigating vascular mechanisms relevant to human disease.

Although congenic and consomic strains are effective for narrowing down genomic regions of interest, they often encompass multiple genes. To achieve functional validation of specific genetic variants, the field has transitioned toward direct genome editing. Advances in genomic tools, including CRISPR/Cas9 [17, 18], zinc-finger nucleases (ZFNs) [19–22], and the Sleeping Beauty transposon system [23], have further expanded the utility of rat models by enabling precise genetic modification and enhancing their relevance to human disease. Notable examples of these precision models include the SS.*Cyp4a1* strain [10], created using the Sleeping Beauty transposon system to study 20-HETE in SS hypertension. The SD.*Add3* KO, generated via ZFN-mediated targeting, to dissect the role of *Add3* in cerebral and renal function [9, 24]. The *Lrat*^−/−^ rat was developed with CRISPR/Cas9 on a BN background to model human retinal dystrophies [25]. The TgF344-AD rat, a transgenic model expressing human APPsw and PS1ΔE9 on a F344 background, enables the study of how cerebrovascular remodeling and myogenic dysfunction contribute to Alzheimer’s disease (AD) progression [26, 27]. Together, these genomic resources provide a powerful framework for studying a wide range of biological processes across multiple physiological systems. By enabling precise genetic manipulation, these models facilitate the study of intricate phenotypes, including the myogenic response, across diverse disease states.

The middle cerebral artery (MCA), the largest branch of the internal carotid artery in both rats and humans, supplies a significant portion of the cerebral hemispheres, including the cortex and basal ganglia [28]. Accounting for approximately 60–80% of the total blood flow from the internal carotid artery and following a relatively direct path, the MCA is the most frequent site of ischemic stroke in humans. Positioned at the junction between high-pressure systemic circulation and low-pressure cerebral capillary beds, the MCA plays a pivotal role in regulating cerebral perfusion. The MCA is essential for cerebral blood flow (CBF) autoregulation via the myogenic response, in which it constricts in response to increased intraluminal pressure. The MCA (along with downstream resistance arterioles) stabilizes CBF, thereby protecting the brain from fluctuations in systemic blood pressure. In Alzheimer’s disease and Alzheimer’s disease-related dementias (AD/ADRD), impaired myogenic response and CBF autoregulation are associated with blood-brain barrier (BBB) leakage and glial activation. These dysfunctions diminish cerebral perfusion and enhance neuroinflammation, contributing to neurodegeneration and ultimately leading to cognitive decline [29].

Because MCA dysfunction and structural remodeling drive the progression of stroke and vascular dementia, characterizing these physiological mechanisms is essential for developing therapeutic and lifestyle interventions. While pressure myography of cerebral arteries has been described previously, existing protocols lack detailed anatomical guidance for consistent MCA M2 segment identification across diverse rat strains. We present this protocol to provide a precision methodology for isolating and studying the structure and function of the rat MCA *ex vivo*. This chapter details the transition from micro-dissection to steady-state pressure myography, allowing for the precise quantification of myogenic reactivity and mechanical properties independent of systemic influences. This methodology bridges the gap between molecular genetic findings and the fundamental physiological control of cerebral circulation. The following protocol has been optimized and validated across diverse rat genetic models, including TgF344-AD [13, 27, 30] T2DN [13, 14, 31, 32], FHH.1^BN^ [33, 34], SS.5^BN^ [10], and other genetically modified strains[8, 10, 33, 35, 36] with technical details in the Notes section.

## 2. Materials

### 2.1. Solutions

1. Physiological Salt Solution (PSS_Ca_): 119 mM NaCl, 4.7 mM KCl, 1.2 mM MgSO_4_, 1.2 mM NaH_2_PO_4_,18 mM NaHCO_3_, 10 mM glucose, 5 mM HEPES, 1.6 mM CaCl_2_. Use ultra-pure water and adjust the pH to 7.4 by bubbling with 95% O_2_ and 5% CO_2_ or a normoxic gas mixture of 21% O_2_, 5% CO_2_, and 74% N_2_ (**Note 1**).
2. Calcium -free physiological salt solution (PSS_0Ca_): 119 mM NaCl, 4.7 mM KCl, 1.2 mM MgSO_4_, 1.2 mM NaH_2_PO_4_, 18 mM NaHCO_3_, 10 mM glucose, 5 mM HEPES, 0.03 mM EDTA. Use ultra-pure water and adjust the pH to 7.4 by bubbling with 95% O_2_ and 5% CO_2_ or a normoxic gas mixture of 21% O_2_, 5% CO_2_, and 74% N_2_ (**Note 1**).
3. Dissection solution: 0.5 - 1% (w/v) bovine serum albumin (BSA) in PSS₀_Ca_. Prepare a 10% (w/v) BSA stock solution by dissolving lyophilized BSA powder in PSS₀_Ca_, then dilute to a final concentration of 0.5 - 1% in PSS₀_Ca_. Stir at room temperature for 1 hour to ensure complete dissolution, then place on ice for use (**Note 2**).
4. Potassium Solution: To prepare 60 mM KCl PSS_Ca_, modify the standard buffer by increasing the KCl concentration to 60 mM and equimolarly reducing NaCl to 63.7 mM to maintain the physiological osmolarity of approximately 275-295 mOsm/L. Adjust the final solution to pH 7.4 by bubbling with 95% O_2_ and 5% CO_2_, or with a normoxic gas mixture of 21% O_2_, 5% CO_2_, and 74% N_2_ (**Notes 1, 3**).

### 2.2. Microsurgical and Dissection Tools (Figure 1)

1. Surgical instruments: Standard dissection scissors, bone cutting forceps, straight forceps with fine tips, and Vannas spring scissors with a straight cutting edge (**Note 4**).
2. Dissecting dishes: Glass dishes with a black silicone elastomer base (**Note 5**).
3. Insect pins: Extra-fine stainless-steel pins for securing the brain or large vessels during microdissection (**Note 6**).
4. Stereomicroscope (10x to 80x magnification): High-resolution dissecting microscope (binocular or trinocular) connected to a weighted base and equipped with a cold-light LED source (**Note 7**).
5. Nylon thread: Multi-filament twisted nylon thread specifically selected for tying blood vessels to a glass cannula (**Note 8**).

### 2.3. Cannula Preparation (Figure 2)

1. Cannula: Borosilicate glass micropipettes (inner diameter: 0.68 mm; outer diameter: 1.2 mm; **Note 9)**.
2. Pipette Puller: A vertical-pull pipette puller equipped with an automated double-mode function (**Note 10**).

### 2.4. Pressure Myography System (Figure 3)

1. Chamber: Self-heated single vessel chamber specifically designed to minimize bath volume for applications involving limited reagents (**Note 11**).
2. Temperature controller. A high-performance temperature controller that is connected to the chamber via a thermistor temperature sensor (**Note 12**).
3. Pressure servo controller with peristaltic pump. A pressure system equipped with a miniature peristaltic pump and a flow-through pressure transducer (**Note 13**). The system must be zeroed before each experiment and the gain adjusted periodically to ensure pressure accuracy (**Note 14**).
4. Pressure transducer: A high-quality, flow-through pressure transducer built with luer-style connector ports for easy integration into the perfusion path (**Note 15**).
5. Gas tank and gas line: A gas supply system used to maintain physiological pH and oxygenation levels within the experimental solutions (**Notes 1 and 16**).
6. A high-resolution trinocular inverted microscope equipped with a digital camera and a series of objectives (4x, 10x, 20x, and 40x; **Note 17**). The imaging system for diameter analysis needs to be calibrated every 3-6 months (**Note 18**).
7. Data Acquisition System: A desktop computer loaded with AmLite software for real-time monitoring and data recording (**Note 19**).

## 3. Methods

### 3.1. Isolation of Middle Cerebral Artery (Figure 4)

1. Rats are anesthetized via isoflurane inhalation (4% induction dose in 100% O_2_ for 2–3 minutes) until there is no toe response to stimulation. Following deep anesthesia, rats are humanely euthanized by rapid decapitation using a calibrated, sharp stainless-steel guillotine in strict accordance with IACUC-approved guidelines (**Note 20**).
2. The head is immediately placed in a petri dish on ice.
3. Utilizing the bone cutting forceps, begin at the foramen magnum and take small, controlled bites of the skull, moving anteriorly along the temporal-parietal sutures on both sides to accommodate the thickness of the rat calvarium. Once the lateral bone channels are established, insert the flatter, slightly curved end of the spatula into a lateral channel and use it to gently leverage and peel the parietal and frontal plates upward and away from the brain, avoiding any compression of the underlying tissue. With the brain dorsal surface exposed, use the tips of the dissection scissors to sever the olfactory bulbs and optic nerves, then insert the narrow, flatter tip of the spatula beneath the ventral surface of the brain to lift it cleanly from the cranial vault and transfer it into the cold dissection dish. Any excess blood is washed away with cold dissection solution (**Note 21**).
4. An approximately 5 x 3 mm cortical section containing the MCA is carefully dissected from the ventral surface of the brain. The tissue is immediately transferred to a dissection chamber containing ice-dissection solution (**Note 22**).
5. Under a stereomicroscope, the pia mater is gently separated from the underlying tissue to expose the vascular structure (**Note 23**).
6. The exposed vasculature is inspected to identify branch-free M2 segments of the MCA. Ideal segments for cannulation are those with diameters ranging from 100 to 180 μm, with a uniform appearance and free of visible side branches or blood clots (**Note 24**).
7. The identified M2 segments are carefully excised using Vannas spring scissors and transferred to a petri dish containing fresh, ice-cold PSS_0Ca_ and maintained on ice for the subsequent cannulation (**Note 25**).

### 3.2. Pressure Myography (Figure 3)

1. Secure the glass micropipettes into the cannula holders within the pressure myography chamber. Align the tips by adjusting the Up & Down and Forward & Backward micromanipulators until they are perfectly centered (**Note 26**).
2. Flush the cannulas with dissection solution containing 0.5 - 1% BSA to prevent vessel adhesion. Subsequently, rinse and fill the cannulas with pre-warmed PSS_Ca_ (**Note 27**).
3. Fill the bath with 6 mL of PSS_Ca_. Place two loose loops of monofilament nylon thread over each cannula tip (**Note 28**).
4. Carefully transfer the dissected MCA into the chamber bath (**Note 29**).
5. Using fine-tipped forceps, gently open the vessel lumen. Use fine-tip forceps to stabilize the vessel edge while carefully sliding the MCA onto the first cannula. Secure it by tightening the two pre-made nylon loops. Repeat this process to secure the distal end of the vessel to the second cannula (**Note 30**).
6. Transfer the loaded chamber to the microscope stage. Connect the temperature probe, the gas line, and the pressure controller connected with the pressure transducer (**Note 31**).
7. Launch the data acquisition software to begin real-time monitoring of the internal diameter (**Note 32**).
8. Gradually increase the intraluminal pressure of the MCA to 40 mmHg. Equilibrate the vessel at 37°C for approximately 30 minutes to restore metabolic activity and establish the ionic gradients required for the development of spontaneous myogenic tone (**Note 33**).
9. Assess vessel viability by monitoring contractile responsiveness during a brief exposure to 60 mM KCl. A viable MCA should exhibit ∼ 25 % constriction; if the vessel fails to meet the 25% threshold, discard and cannulate a new vessel (**Note 34**).
10. Replace the bath with warm PSS_Ca_ and allow the vessel to stabilize for 3 - 5 minutes at 40 mmHg. Gradually increase the intraluminal pressure in 20 mmHg increments, maintaining each level for 3 - 5 minutes (or until a steady state is reached) up to 140 - 180 mmHg, depending on the strain (**Note 35**).
11. Change the bath solution to warm PSS_0Ca_ and stabilize the vessel at 40 mmHg. Perform a stepwise pressure ramp by increasing the intraluminal pressure in 20 mmHg increments to a maximum of 140-180 mmHg, as determined by strain (**Note 36**).
12. Following the completion of the passive pressure-diameter curve, use the recorded inner diameter (*ID_oCa_*) and outer diameter (*OD_oCa_*) to calculate the following structural and mechanical parameters.

◦ Wall Thickness (WT): 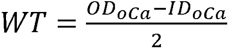
◦ Wall-to-Lumen Ratio: 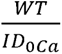
◦ Cross-Sectional Area (CSA): 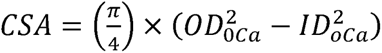
◦ Myogenic Tone (%): 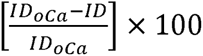
◦ Distensibility (%): 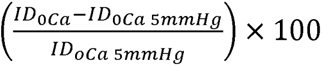(**Note 37**)
◦ Incremental Distensibility (% /mmHg): 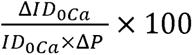(**Note 38**)
◦ Circumferential Wall Strain (ε): 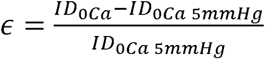
◦ Circumferential Wall Stress (σ): 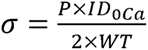
◦ Elastic Modulus (β value): Determine this by fitting stress-strain data to the exponential model: *σ* = *αe^β∈^*

- *α*: The intercept (theoretical stress at zero strain)
- *β*: The slope of the exponential fit; a higher β value indicates increased arterial stiffness.

## 4. Examples of Expected Results

Examples of expected results are presented in **Figure 5**.

**Figure 1.**
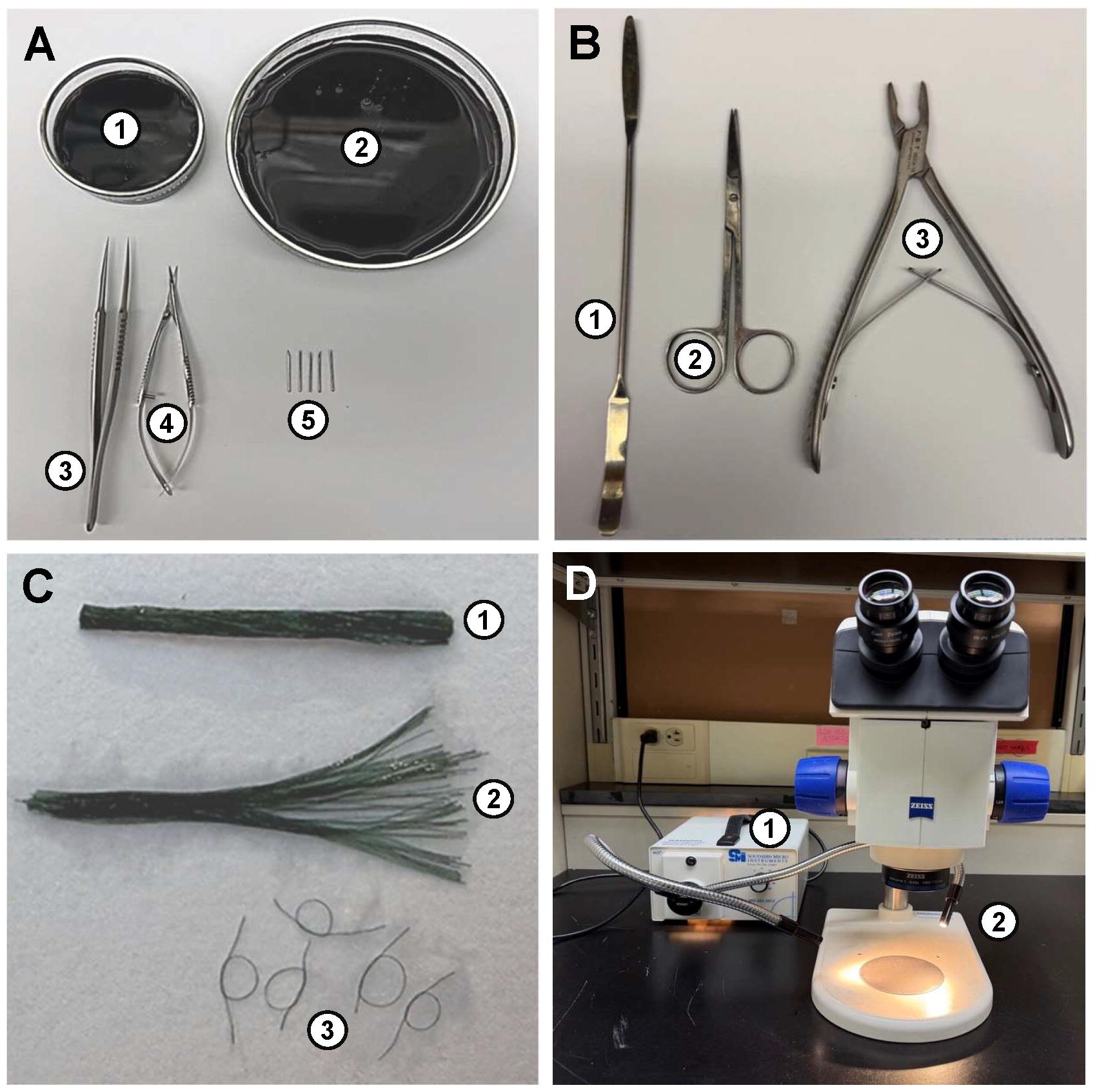
Microsurgical and Dissection Tools. **A:** Fine dissection tools and dishes. (**1, 2**) Glass dissecting dishes of varying sizes containing a black silicone elastomer base. (**3**) Straight forceps with fine tips. (**4**) Vannas spring scissors with a straight cutting edge. (**5**) Extra-fine stainless-steel insect pins. **B:** Gross surgical instruments. (**1**) Spatula for brain removal. (**2**) Standard dissection scissors. (**3**) Bone cutting forceps. **C:** Suture. (**1**) Multi-filament twisted nylon thread (∼ 500 μm), which is separated into individual 30 μm monofilaments (**2**). (**3**) Individual loops made from monofilaments. **D:** Stereomicroscope setup. High-resolution binocular dissecting microscope equipped with (**1**) a cold-light LED source and (**2**) dual gooseneck fiber optic illuminators to provide adjustable, non-thermal lighting during microdissection.

**Figure 2.**
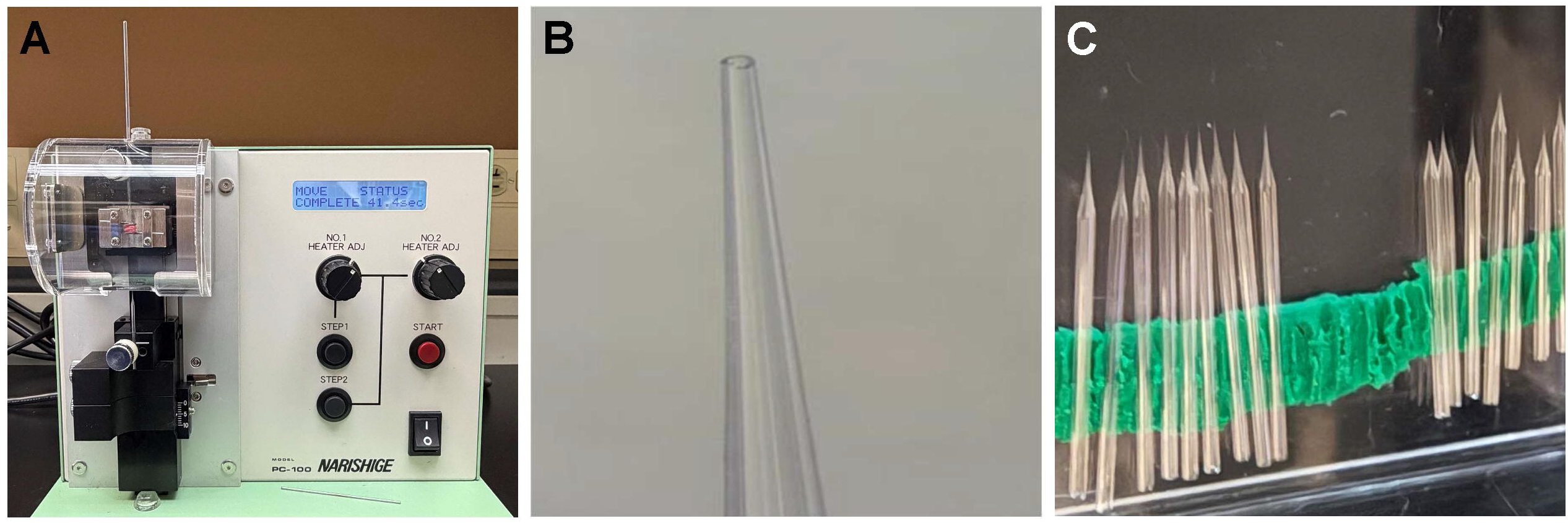
Micropipette Fabrication for Rat MCA Cannulation. **A:** A vertical micropipette puller used to produce glass micropipettes from borosilicate capillaries. Parameters are optimized via a two-stage pull to ensure consistent taper geometry. **B:** Microscopic view of a pulled micropipette that is precision-broken and fire-polished to an outer diameter of ∼80 μm. **C:** Pulled pipettes are secured on a sculpting clay base within a covered container for storage.

**Figure 3.**
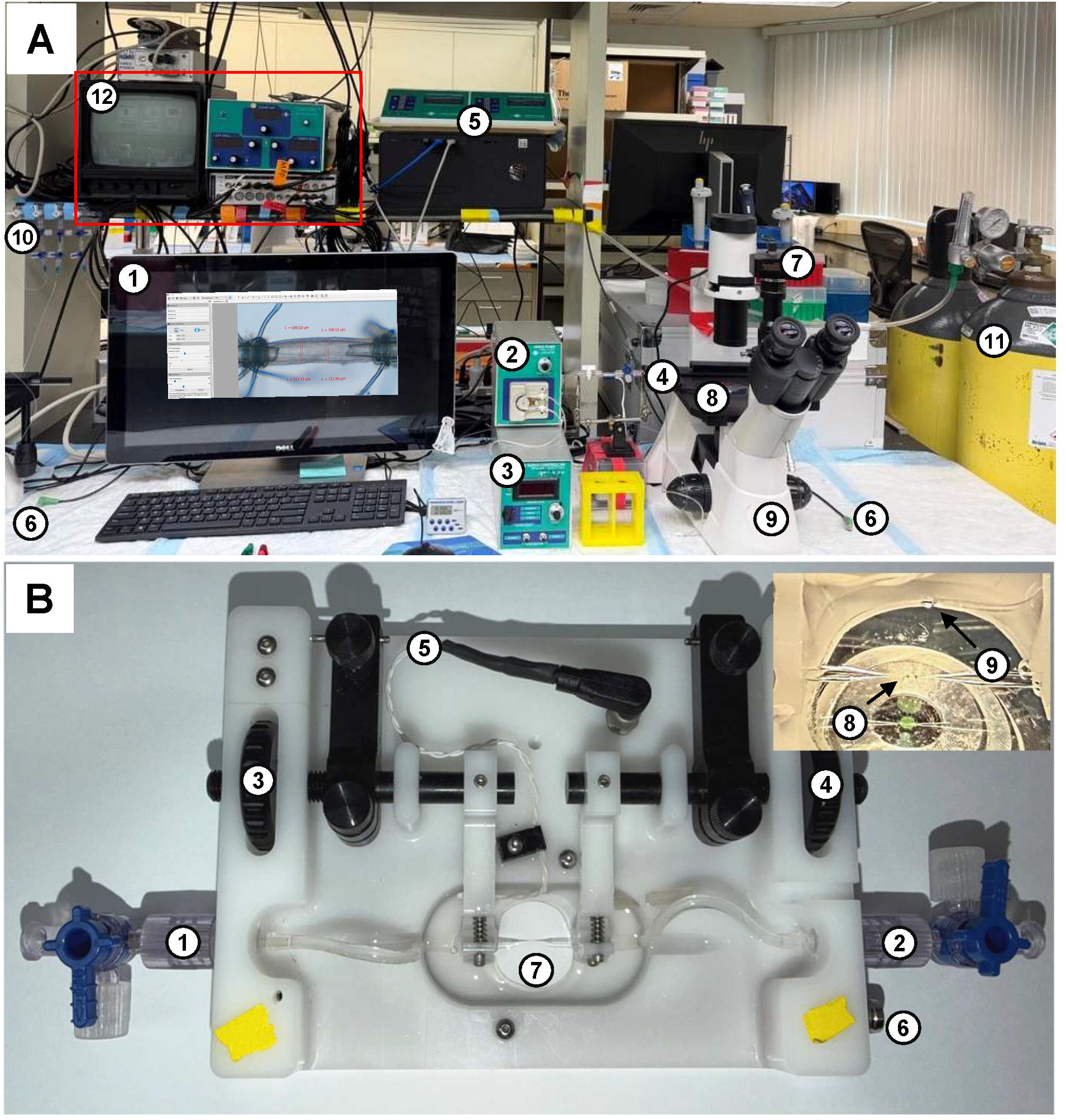
Pressure Myography Workstation and Chamber. **A:** Pressure myography workstation. (**1**) Data acquisition system: Desktop computer interface for real-time diameter monitoring and data recording. Insert: an example of a mounted rat MCA as captured by the data acquisition software for real-time diameter tracking and edge detection. (**2, 3**) Pressure servo controller and peristaltic pump. (**4**) Pressure transducer. (**5**) Temperature controller. (**6**) Thermistor probes. (**7**) Digital camera connected to the microscope. (**8**) The stage-mounted platform where the cannulated vessel is placed. (**9**) Inverted trinocular microscope. (**10**) Gas manifold: three-way valves for distributing gas to multiple stations. (**11**) Gas cylinders. (**12**) Alternative data acquisition system including a monitor and a video dimension analyzer integrated with a PowerLab interface and a pulse generator amplifier. **B:** Pressure myography chamber. (**1, 2**) Inflow (pressure) and outflow ports. (**3, 4**) Micromanipulators: Precision knobs used for the lateral alignment of micropipettes and for adjusting the longitudinal (axial) stretch of the vessel to its *in vivo* length. (**5, 6**) Thermistor and electrode connecting to the temperature controller. (**7**) Central glass viewing port with aligned micropipettes mounted, ready for vessel cannulation. Insert: (**8**) The arrow indicates a cannulated vessel secured between aligned micropipettes with monofilament ties. (**9**) The arrow indicates a gas line providing continuous bubbling of carbogen to maintain physiological pH and oxygenation without disturbing the vessel position or interfering with the optical clarity of the real-time diameter tracking.

**Figure 4.**
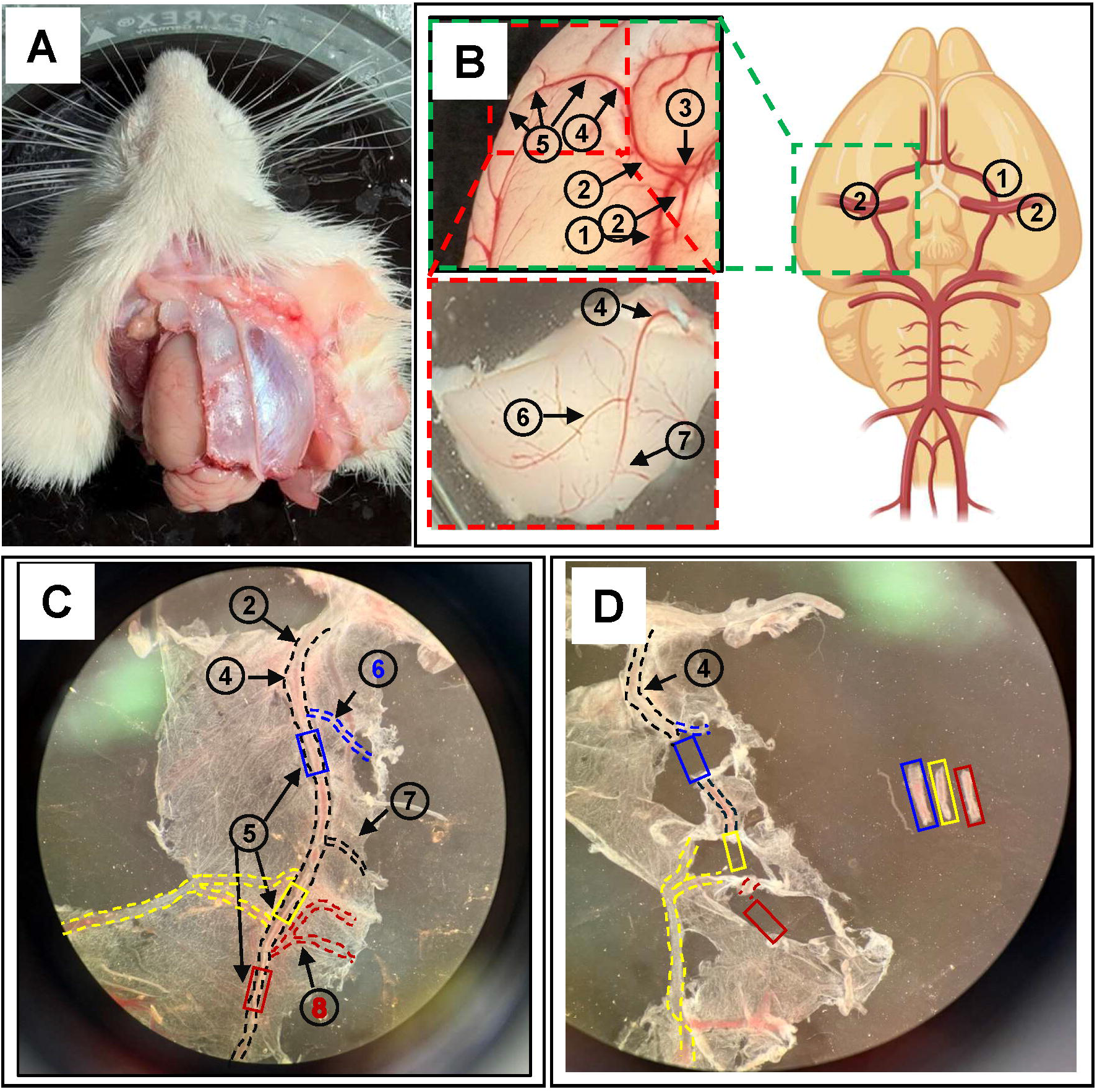
Anatomical Identification and Microdissection of the Rat Middle Cerebral Artery (MCA). **A:** Ventral view of the rat head post-decapitation, demonstrating the initial surgical exposure required for brain removal and access to the cerebral vasculature. **B: *Right:*** Schematic illustration of the Circle of Willis (**1**) and the bilateral distribution of the MCA, generated by BioRender. ***Left:*** *In situ* view of the cerebral vasculature showing the MCA M1 segment (**2**) and the adjacent rhinal vein (**3**). The “Turn” (**4**) marks the critical anatomical landmark where the vessel transitions from the ventral surface to the lateral surface, becoming the MCA M2 segment (**5**). The bottom-left inset shows a 3 x 5 cm section of the surgical field to show the further distal end of the M2 segment. **C:** High-magnification microscopic view of the isolated pial membrane containing the MCA tree. This view captures the MCA M1 segment (**2**) before the “Turn” (**4**) and its transition into the MCA M2 segment (**5**). The branching pattern of the M2 is clearly visible, leading to downstream bifurcations. Colored dashed boxes indicate the specific segments selected for microdissection: blue: post 1^st^ (**6**), bifurcation; yellow: post 3^rd^ bifurcation; and red: post 5^th^ bifurcation (**8**). **D:** Final stage of microdissection with MCA segments cleared of connective tissue and separated. The isolated segments are indicated by the color-coded rectangles described in C. In general, M2 segments located post the “Turn” or within the first two bifurcations (diameters 100-180 µm) are optimal for cannulation, with distal segments exhibiting progressively smaller diameters. An isolated penetrating arteriole downstream of the M1 is positioned to the left for comparison. Legend Key: (**1**) Circle of Willis. (**2**) MCA M1 segment. (**3**) Rhinal vein. (**4**) The “Turn” (M1 to M2 transition). (**5**) MCA M2 segments. (**6**) 1^st^ Bifurcation of MCA M2. (**7**) 2^nd^ bifurcation of MCA M2. (**8**) 5^th^ bifurcation of MCA M2.

**Figure 5.**
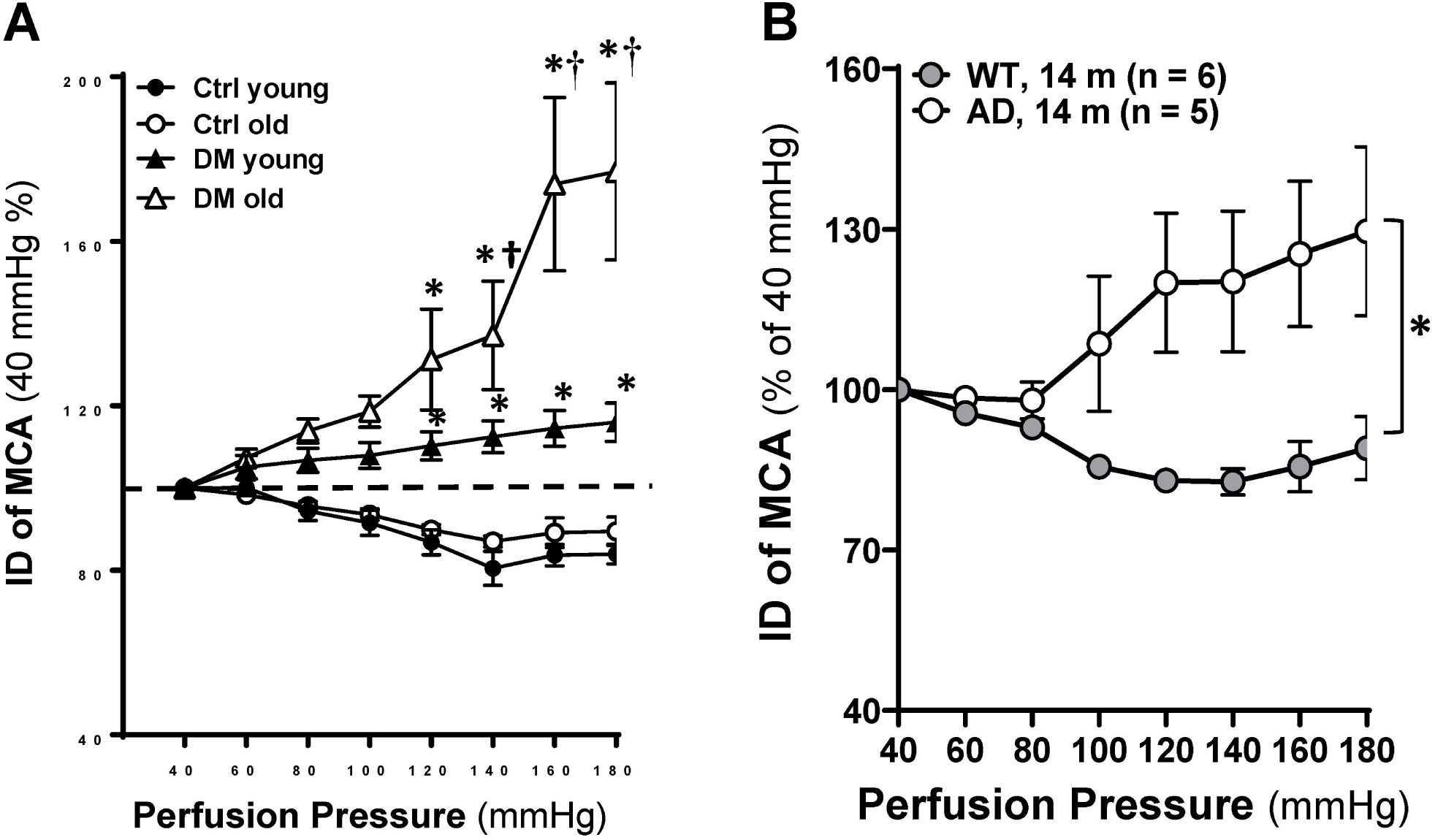
Representative Pressure-dependent Responses of the Rat Middle Cerebral Artery (MCA) Measured by Pressure Myography. **A:** Representative pressure-diameter relationship from 3- and 18-month-old diabetic (DM) T2DN rats and non-diabetic Sprague Dawley (SD) controls. Data from *Wang et al.* [14], reproduced with permission. **B:** Representative pressure-diameter relationship from 14-month-old TgF344-AD rats and age-matched F344 controls. Data from *Fang et al.* [27], reproduced with permission.

Young SD rats (3 months) display an intact myogenic response in the MCA (**Figure 5A**). As intraluminal pressure is raised from 40 to 140 mmHg, the vessel undergoes progressive, active vasoconstriction; the diameter typically stabilizes after ∼3–5 min at each step. This pattern matches the three⍰phase model of myogenic behavior: phase 1: development of myogenic tone at 40 mmHg; phase 2: active myogenic response at around 60-140 mmHg; phase 3: high⍰pressure stage in which autoregulatory breakthrough occurs and active constriction is lost, producing forced (passive) dilation, typically at 160-180 mmHg in the healthy strain and seen as a modest increase in diameter relative to peak constriction [14, 29]. Advanced age alone (e.g., 18 months) does not markedly alter this pattern in SD rats: the pressure-diameter relationship and high⍰pressure breakthrough remain broadly similar, indicating preserved pressure-induced cerebrovascular reactivity in this healthy strain. A similar pattern has also been observed in F344 rats [27].

In contrast to the intact myogenic response in SD rats, T2DN/diabetic (DM) animals exhibit impaired myogenic reactivity. T2DN/DM rats serve as a model of diabetes⍰associated ADRD (DM⍰ADRD) to investigate how chronic metabolic disease drives cerebrovascular dysfunction and dementia [13, 14, 32, 37, 38]. This impairment is already evident at 3 months of age and exacerbates progressively with aging. In elderly DM rats, the myogenic response shows a severe, pressure-dependent loss, causing the upper autoregulatory breakthrough point to shift to the left, with forced dilatation becoming highly apparent at pressures above 140 mmHg (**Figure 5A**) [14]. A similar pattern of impaired myogenic response in the MCA is also reported in TgF344-AD rats, an established model of AD [30, 39, 40]. Longitudinal characterization of cerebral hemodynamics in these AD rats reveals that myogenic responses in cerebral arteries and arterioles become compromised at 4 months of age, two months prior to the onset of cognitive decline [27]. Consistent with this *ex vivo* finding, the AD model exhibits poor autoregulation, and this cerebrovascular hemodynamic dysfunction is significantly exacerbated with advanced age; by 14 months of age, the phenotype transitions to a critical, severe pattern of reduced cerebral perfusion and profound autoregulatory failure (**Figure 5B**) [27].

Together, these examples demonstrate the expected range of physiological and pathological phenotypes attainable with this protocol, providing critical benchmarks for validating vessel viability and evaluating experimental success.

## 5. Notes

1. The pH is maintained at 7.4 by continuous bubbling with a gas mixture of 95% O_2_ and 5% CO_2._ Since the vessel is removed from its natural blood supply, it relies entirely on dissolved oxygen in the buffer, and a higher partial pressure of oxygen (pO_2_) may be needed to ensure oxygen diffuses through the vessel wall. However, the physiological normoxic gas mixture (21% O_2_, 5% CO_2_, and 74% N_2_) is preferred to minimize hyperoxia-induced oxidative stress, provided the vessel remains viable throughout the experiment.
2. BSA is added to the dissection buffer to reduce surface tension, limit non-specific protein binding, and prevent the delicate MCA from adhering to the glass dish or microsurgical tools. It is critical to use lyophilized (powdered) BSA, such as Fraction V or fatty acid-free grade, and dissolve it directly into the prepared PSS₀_Ca_. Commercially available BSA stock solutions prepared in Phosphate-Buffered Saline (PBS) must be avoided, as they introduce exogenous sodium, chloride, and phosphate ions. These additions disrupt the precise ionic balance and osmolarity of the physiological salt solutions, which can significantly impair the myogenic response. Furthermore, many liquid BSA stocks contain stabilizers or trace amounts of EDTA that can interfere with calcium-dependent vascular tone, even at very low concentrations.
3. For the rapid viability check, use the iso-osmotic 60 mM KCl PSS_Ca_. Exchanging the bath volume 5-6 times ensures the target concentration is reached while keeping the vessel safely submerged and free of mechanical stress.
4. Our lab typically uses straight forceps for dissection and mounting steps, but curved versions are also effective depending on the depth of the surgical field and the specific angle required for precise tissue manipulation.
5. The black silicone base in the dissection dishes provides visual contrast for the white brain tissue and a surface for pinning. Deep or shallow dishes may be used depending on the volume of dissection buffer required to keep the tissue submerged and the working distance available under your microscope objective.
6. Alternatively, 26 or 28-gauge needles can be used to secure delicate tissue to the silicone elastomer base for precise dissection.
7. Using a binocular or trinocular stereomicroscope provides the depth perception necessary for precise vessel isolation, while a weighted base, some paired with a boom arm, ensures structural stability and sufficient working space. Additionally, an LED cold-light source is essential for maintaining the physiological integrity of the tissue by preventing heat transfer and buffer evaporation during dissection. The dissecting microscope we are using features 10x eyepieces and an adjustable zoom magnification range of 1.0 to 8.0.
8. Nylon thread is often used due to its consistency and is less likely to cause a reaction or interact with sensitive biochemical solutions [41, 42]. Our lab uses a thread with an overall diameter of approximately 500 μm, composed of individual monofilaments roughly 30 μm in diameter. To use, we tease away and tear off a single monofilament to create the overhand knots required to secure the vessel to the cannula.
9. Borosilicate glass provides the close dimensional tolerances required for uniform, reproducible microelectrodes.
10. The vertical pipette puller our lab uses features an automated double-mode function and utilizes gravity-fed pulling forces. To generate cannulas with long, tapered tips (∼80 μm), install two “light type” weights (Variation 2) and select the two-stage pull mode. Configure the puller with a Preheating level of 60.0, a Heater level of 100 (2.5 V) for both stages, and durations of 15 s for Step 1 and 10 s for Step 2.
11. This system is integrated with a temperature controller and a thermistor temperature sensor. This configuration allows for the vessel chamber to be heated directly to physiological temperatures, ensuring stable thermal conditions without the need for constant bath superfusion.
12. Our lab uses a temperature controller featuring a novel control algorithm that minimizes the time to reach physiological setpoints (typically 6–10 min from 22°C to 37°C).
13. The system operates in either pressure mode (0–200 mmHg) to maintain a constant transmural set-point or flow mode to provide a stable perfusion rate. When configured with the pressure pump, a flow-through pressure transducer is needed to provide the continuous feedback required for precise regulation of intravascular pressure.
14. The calibration process involves two distinct steps: Zeroing (required prior to each experiment) and Gain Calibration (recommended on a monthly basis or whenever transducer sensitivity verification is required). To zero the system, open the transducer to atmospheric pressure and rotate the ZERO dial on the front panel until the display reads 0 mmHg. To calibrate the gain, apply a known pressure (e.g., 100 mmHg) using a manometer and adjust the GAIN screw on the back panel until the display matches the manometer reading.
15. The pressure transducer features a range of - 50 to 300 mmHg that is compatible with the pressure servo controller.
16. The gas tank is connected to the system using polyethylene (PE) tubing. This line is directed into the experimental chamber to ensure the physiological salt solutions remain appropriately buffered and oxygenated. For setups with multiple stations, a 3-way controller is used to distribute the gas supply efficiently among the chambers.
17. The trinocular head of the microscope is connected to a digital camera, which transmits the live video feed to the data acquisition system for diameter analysis. A vibration isolation table could be used to minimize mechanical noise.
18. To calibrate the imaging system for diameter analysis, place a certified stage micrometer on the microscope stage and bring the scale into sharp focus using the same objective and zoom settings required for the MCA experiment. Using data acquisition software, capture a high-resolution image of the micrometer and draw a reference line across a known physical distance (typically 500 μm or 1 mm) to map the number of pixels to micrometers. Save this calculated calibration factor as a dedicated profile, and verify its accuracy by measuring a separate known interval on the scale, ensuring the vibration-isolation table remains active throughout the process to prevent mechanical noise from distorting the pixel-to-micron conversion.
19. AmLite software is utilized in our laboratory and installed on a dedicated desktop computer to facilitate high-speed data acquisition. This software interface enables simultaneous tracking of vessel diameter, intravascular pressure, and temperature across multiple stations, providing a comprehensive digital record of the experimental parameters. Alternatively, an analog-to-digital data acquisition system, combined with a Video Dimension Analyzer and an external monitor, may be used to consolidate pressure, temperature, and diameter signals into a single, synchronized digital record [41].
20. The guillotine must be cleaned with 70% ethanol, and the blades are inspected prior to each session to ensure a swift, one-stroke procedure.
21. To prevent cortical penetration or microvascular damage, maintain the bone cutting forceps parallel to the skull and ensure the Circle of Willis remains intact; use the spatula for gentle leverage rather than downward pressure, completing the entire process from decapitation to immersion within 2-3 minutes to preserve the metabolic integrity and myogenic reactivity of the middle cerebral arteries.
22. To dissect the cortical section in rats, first identify the Circle of Willis on the ventral surface to locate the MCA as it emerges laterally from the internal carotid artery. Focus the tissue block specifically on the area where the MCA passes dorsal to the olfactory tract and enters the rhinal fissure. Ensure the primary M1 and M2 branches are centered within the block to simplify identifying and removing the pia mater in subsequent steps.
23. To stabilize the tissue for precise dissection, use stainless steel insect pins to secure the edges of the cortical section to the black silicone elastomer base of the dissection chamber.
24. To locate the M2 (insular/sylvian) segment of the rat MCA, follow the M1 (sphenoidal) trunk from the Circle of Willis toward the lateral surface of the brain. The key landmark for the M2 transition is the 90-degree “turn” where the artery crosses both the rhinal fissure and the rhinal vein. At this junction, the vessel moves from the base of the brain onto the superficial pial surface of the piriform cortex. Select a straight, branch-free segment immediately distal to the “Turn”, which could be proximal or distal to the first or second major bifurcation (typically the frontal or parietal branches). Ensure the target vessel remains on the pial surface and does not give off any deep lenticulostriate perforators. For consistent experimental results, prioritize a segment with a uniform diameter of 100-180 µm and a usable length of 1.5-2.0 mm.
25. During dissection, use fine micro-pins to secure the surrounding pia mater or the connective tissue near the vessel edges; this provides the necessary tension to isolate the segment without directly pinning the vessel wall itself. When transferring, gently grasp only the extreme edge of the vessel with fine-tipped forceps to avoid mechanical damage to the experimental zone. The segments should ideally be kept in cold PSS_0Ca_ and used within 4-6 hours to maintain optimal physiological responsiveness.
26. Proper alignment is critical to ensure the vessel remains straight and untwisted. Misalignment can create uneven wall tension, leading to asymmetrical myogenic responses or premature vessel rupture during pressurization.
27. BSA prevents the MCA from sticking or tearing during mounting. The use of PSS_Ca_ is strictly required to provide the extracellular calcium necessary for the development of myogenic response. While cannulation can be performed at room temperature for easier handling, the system must eventually be warmed to 37°C to initiate physiological activity.
28. Ensure the monofilament loops are large enough to slide easily over the cannula but small enough to be precisely positioned with fine-tipped forceps. Keeping the loops near the tip of the micropipette will facilitate rapid securing of the vessel ends during the cannulation step.
29. Use fine-tipped forceps to grasp only the very end of the vessel or surrounding connective tissue to avoid damaging the functional trunk. Ensure the vessel remains submerged during the move to prevent lumen collapse or mechanical trauma caused by surface tension.
30. Maintain a small gap between the pipette tip and the first tie to ensure the lumen remains unobstructed. Avoid over-stretching the MCA when connecting the distal end, as excessive longitudinal tension can alter the vessel’s pressure-diameter relationship. Confirm that the knots are tight enough to prevent leaks without cutting through the vessel wall.
31. Confirm that all plumbing connections are airtight and that the pressure transducer is leveled with the vessel to avoid hydrostatic pressure errors. Pre-zeroing the system before mounting is essential to ensure that the reported intraluminal pressure is accurate. Set the pressure controller to a low initial pressure (e.g., 5 mmHg) to maintain vessel patency without over-stretching the tissue. Adjust the gas flow to a level that ensures proper aeration without creating excessive turbulence that might vibrate the vessel.
32. Select a clean section of the vessel for the tracking zone, ideally away from the cannula tips or any remaining connective tissue. Once tracking is established, verify that the software is correctly identifying both the inner and outer walls. Consistent monitoring at this stage allows for the early detection of leaks or loss of vessel viability before the experiment proceeds. To ensure data integrity, maintain constant camera settings (exposure, gain, and light intensity) throughout the recording to prevent imaging artifacts.
33. Successful equilibration is marked by the development of spontaneous myogenic tone. Normal rat strains typically exhibit a robust response, resulting in a ∼30% decrease in diameter from the initial passive state.
34. To avoid damaging surface tension, never empty the chamber. Perform a simultaneous buffer exchange by adding the premixed warm 60 mM KCl in PSS_Ca_ at one end while aspirating from the other. Exchanging 5-6 times the bath volume ensures the target concentration is reached while keeping the vessel safely submerged and free of mechanical stress.
35. Similarly, use a simultaneous buffer exchange method to prevent exposing the vessel to the air-liquid interface. Allow 3-5 minutes at each step (or until a steady state is reached) before recording data. While healthy strains are typically tested up to 160-180 mmHg in intraluminal pressure, some diseased strains, such as TgF344-AD [27, 30], T2DN [13, 14, 31], FHH [33], and SS [43] may require only up to 140-160 mmHg to fully characterize the response, as their autoregulatory breakthrough points are often shifted to lower pressures. Throughout the protocol, measure the inner and outer diameters at a minimum of two consistent points along the vessel, ensuring these measurements are closely aligned and that the same coordinates are used for all subsequent steps to maintain data consistency.
36. Achieving a fully passive state requires the absolute removal of calcium from the bath and tissue. Use the simultaneous buffer exchange method to wash the chamber, which typically requires 6 or more volume exchanges; we include a calcium chelator, EDTA (0.03 mM), in PSS0Ca to ensure that any calcium leaching from the tissue is effectively sequestered. If a standard 3-5-minute stabilization is insufficient for this step, a longer equilibration period is required to ensure the vessel reaches a stable maximum diameter. Ensure the vessel has returned to its initial dimensions before starting the pressure ramp. It is recommended to first establish a full myogenic response curve (40-180 mmHg) to define the autoregulatory limits of the specific strain in pretested rats, without recording passive diameters. Once the vessel has surpassed its autoregulatory breakthrough point, it cannot be used for structural analysis [44].
37. *ID*_0*Ca 5mmHg*_ refers to the *ID_oCa_* obtained at the perfusion pressure of 5 mmHg in PSS_0Ca_.
38. Incremental distensibility is defined as the percentage change of the vascular *ID_oCa_* for every 1 mmHg (= 133.4 *N m*^−2^) change in (*P*), the intraluminal pressure in PSS_0Ca_. ΔP indicates the increment in intraluminal pressure in PSS_0Ca_.

